# The Recombination Landscape of the Khoe-San - the Upper Limits of Recombination Divergence in Humans

**DOI:** 10.1101/2021.11.07.467603

**Authors:** Gerald van Eeden, Caitlin Uren, Evlyn Pless, Mira Mastoras, Gian D. van der Spuy, Gerard Tromp, Brenna M. Henn, Marlo Möller

**Author notes:** Corresponding Author: DSI-NRF Centre of Excellence for Biomedical Tuberculosis Research, South African Medical Research Council Centre for Tuberculosis Research, Division of Molecular Biology and Human Genetics, Faculty of Medicine and Health Sciences, Stellenbosch University, Cape Town, South Africa, Fax number: N/A. **Email Addresses:**.

## Abstract

Recombination maps are important resources for epidemiological and evolutionary analyses, however, there are currently no recombination maps representing any African population outside of those with West African ancestry. We inferred the demographic history for the Nama, an indigenous Khoe-San population of southern Africa, and derived a novel, population-specific recombination map from the whole genome sequencing of 54 Nama individuals. We hypothesized that there are no publicly available recombination maps representative of the Nama, considering the deep population divergence and subsequent isolation of the Khoe-San from other African groups. We showed that the recombination landscape of the Nama does not cluster with any continental groups with publicly available representative recombination maps. Finally, we used selection scans as an example of how fine-scale differences between the Nama recombination map and the combined Phase II HapMap recombination map can impact the outcome of selection scans.

## Introduction

Recombination enables the evolution of complex traits by shuffling novel genetic variants, brought about by mutation, into new combinations with existing alleles from varying genomic origins (Peñalba and Wolf 2020). Due to the evolutionary significance of recombination, many implementations of software packages that infer the recombination rate have been developed. Some of these packages rely on inferring past recombination events by analyzing pedigrees (Halldorsson et al. 2019), by detecting changes in ancestry (Hinch et al. 2011; Wegmann et al. 2011) or by the boundaries of blocks of identity by descent (IBD) (Zhou et al. 2020). These methods require large sample numbers (> 2000) to accurately infer the recombination rate at fine-scales. Other packages use linkage disequilibrium (LD) (Auton and McVean 2007) or derivatives thereof, e.g. summary statistics (Gao et al. 2016), to infer recombination into the very distant past and require fewer individuals to infer recombination at fine-scales. LD-based recombination maps, however, are strongly influenced by past demographic events, e.g. population bottlenecks (Dapper and Payseur 2018). Recombination inference software that is aware of changes in the effective population size (N_e_) of a population, such as pyrho (Spence and Song 2019), can be used to mitigate this effect.

The rate of recombination varies between species (Auton et al. 2012), between populations within species (Graffelman et al. 2007; Spence and Song 2019) and even among individuals (Pratto et al. 2014). The recombination rate across the genome is generally expressed as a ratio of genetic distance and physical distance, known as a recombination map. It has been shown that at low resolutions (> 1 Mb), population specific recombination maps are fairly similar (Serre et al. 2005) and at high resolutions they correlate according to continental levels of population differentiation (Graffelman et al. 2007). For instance, the pedigree-based deCODE (Kong et al. 2010) map, based on the Icelandic population, correlates better at fine scales to the linkage-disequilibrium-based (LD-based) HapMap II (International HapMap Consortium et al. 2007) map of the CEU (Utah residents with Northern and Western European ancestry from the CEPH collection) than it does to the HapMap II map of the YRI (Yoruba in Ibadan, Nigeria) (Kong et al. 2010; Gao et al. 2016). Many population-specific recombination maps have been inferred to date, but none have been inferred for any southern African populations (Swart et al. 2020) and researchers studying these populations have had to use available maps that might not suit their analysis.

In this manuscript we present a novel recombination map for the Nama - an indigenous population of southern Africa (Uren et al. 2016) that forms part of a larger group of geographically close and culturally related individuals known collectively as the “Khoe-San”. The Khoe-San are reported to have the most divergent lineages of any other living population (Gronau et al. 2011; Henn et al. 2011; Pickrell et al. 2012; Schlebusch et al. 2012; Barbieri et al. 2016) and it is believed that they have largely remained isolated until ∼2000 years ago (Henn et al. 2011; Barbieri et al. 2013; Uren et al. 2016). Therefore, a recombination map for this population may be very different at fine scales compared to recombination maps that have been inferred for other populations. The Khoe-San also contribute a significant ancestral component (15-75%) to admixed southern African groups, like the South African Coloured (SAC) population and southern Bantu-speaking populations (Uren et al. 2017; Sengupta et al. 2021), and a recombination map for diverse Khoe-San populations could benefit studies involving these groups. The demographic history of the Nama is multi-layered, with 5-25% gene flow from Eastern African caprid and cattle pastoralists ∼2000 years ago (Henn et al. 2008) and genetic exchange with the Damara - a hunter-gatherer population of West-Central African ancestry who became economic clients of the Nama. These events were finally followed by recent admixture with European colonists and to a lesser degree ∼250 years ago.

We used whole genome sequencing (WGS) data of 54 unrelated Nama individuals to infer a LD-based recombination map that is adjusted according to past changes in N_e_. Demographic history was inferred using SMC++ (Terhorst et al. 2017) for distant changes in N_e_ and AS-IBDNe (Browning et al. 2018) for recent N_e_ changes. The N_e_ size changes were then combined and the demography-aware LD-based method pyrho (Spence and Song 2019) was used for recombination rate inference. The resultant population-specific recombination map was then compared to other publicly available recombination maps using the Spearman rank correlation coefficient. Finally, we assessed the fine scale differences between the inferred Nama recombination map and the combined Phase II HapMap recombination map in a region of chromosome 1 and demonstrated how the use of different recombination maps can affect the results from a selection scan.

## Results

### The Inferred Demographic History of the Nama

We inferred the N_e_ for the Nama using SMC++ (Fig. 1 A right) and AS-IBDNe (Fig. 1 A left). The results from SMC++ represent the N_e_ change from 50 000 to 260 generations into the past. The AS-IBDNe results represent the N_e_ change from 50 to 4 generations into the past and the N_e_ was inferred using IBD segments of Khoe-San ancestry exclusively. An N_e_ of ∼30 000 approximately 10 000 generations ago with a reduction in N_e_ to ∼21 000 approximately 5000 generations ago is consistent with previously published inferred N_e_ for the Nama (Schlebusch et al. 2020). Inconsistent with previous results, there is a further reduction in N_e_ to ∼10 000 approximately 1000 generations ago. The inferred N_e_ by SMC++ then stops at 260 generations, because SMC++ can infer N_e_ 6-120 thousand years ago (kya) with low error (Terhorst et al. 2017) and by default SMC++ uses an heuristic to calculate these timepoints automatically given the data.

**Figure 1:**
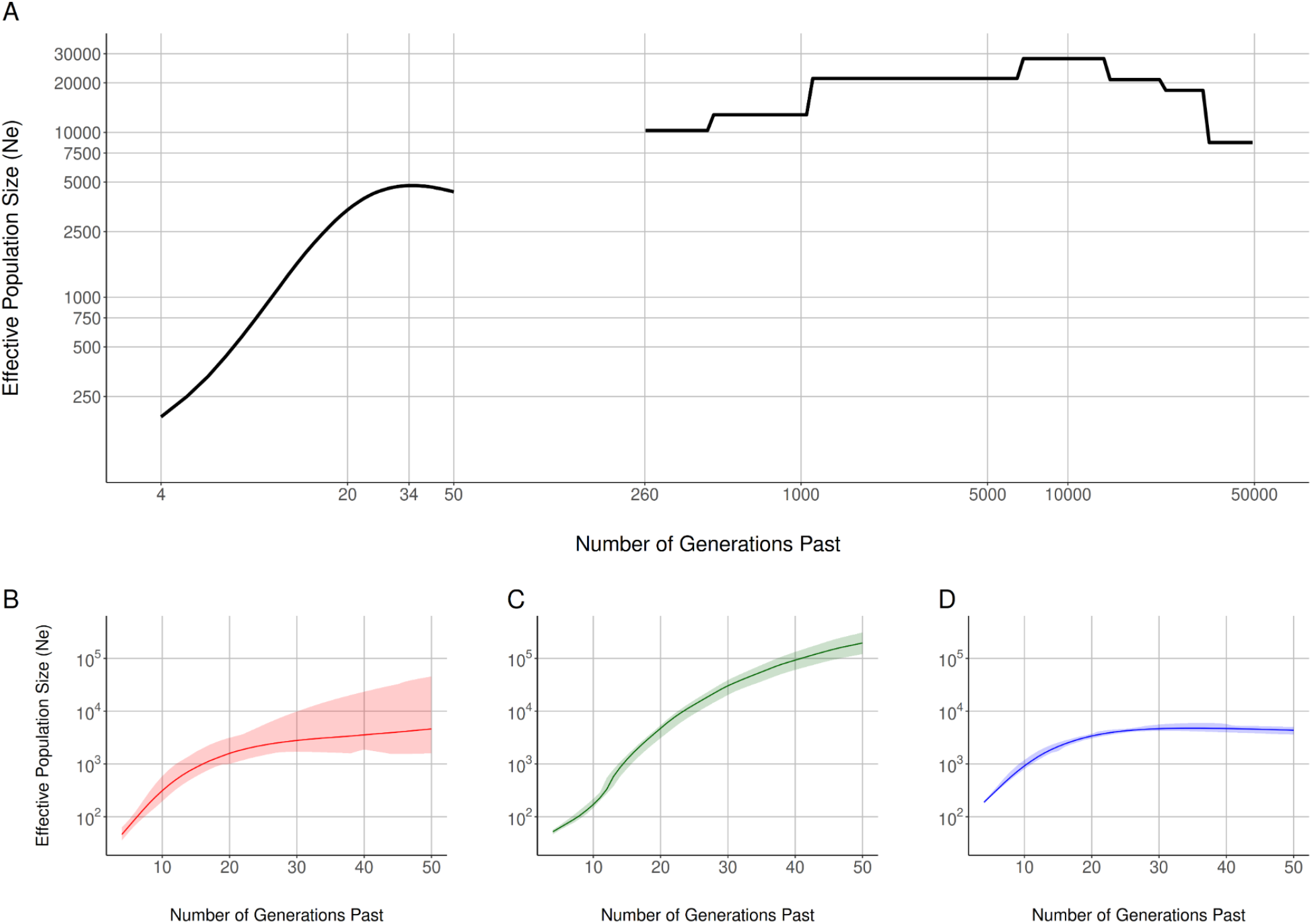
(A) The inferred effective population size history for the Nama plotted on a log10 scale with SMC++ results on the right and AS-IBDNe results for the Nama on the left. (B-D) The AS-IBDNe results for the LWK (B), GBR (C) and Nama (D) ancestral components in the Nama.

We therefore used AS-IBDNe to estimate population fluctuations over the past thousand years (Browning and Browning 2015). We deconvolved 84 Nama genomes (SNP array) into local ancestry tracts with three possible ancestry states: Khoe-San ancestry, European ancestry and Western-Central African ancestry as represented by Nama, GBR (British in England and Scotland) and LWK (Luhya in Webuye, Kenya) population samples. Admixture deconvolution was performed on MEGA SNP array (Martin et al. 2017) data in order to facilitate large numbers of haplotypes in the reference populations. Beginning 50 generations ago, we infer an N_e_ of 4 360. The N_e_ starts to decline 34 generations ago and continues to decline until an N_e_ of 190 inferred 4 generations ago; estimation stops 4 generation ago to avoid coalescent events based on genealogical relationships. The rapid population decline substantially predates the arrival of European settlers in the Richtersveld in 1760 (or ∼7 generations ago), an arid region just south of the Orange River (Smith 1995). The N_e_ results inferred for each set of ancestry specific IBD segments (Fig. 1 B-D) have very narrow 95% confidence intervals.

### The Correlation Between the inferred Nama Recombination Map and Other Publicly Available Maps

Fine-scale recombination rate differences between pairs of populations are correlated according to continental levels of population differentiation (Graffelman et al. 2007; Spence and Song 2019). Considering the long period that the Nama were isolated and their complex demographic history, we hypothesise that there is no available recombination map that is representative of the Nama. Therefore, we compared the inferred recombination map for the Nama with 26 other publicly available recombination maps derived from (Spence and Song 2019) using the Spearman rank correlation at a 2 kilobase resolution (Fig. 2). These maps were inferred for populations from the 1000 Genomes (1000 Genomes Project Consortium et al. 2010) dataset and the populations are classified into various super-populations representing major ancestry differences. We find that pairwise correlations between all 27 maps cluster according to continental levels of population differentiation (super-populations). Furthermore, we find that the Nama are more closely related to other African populations than to other continental groups (< 0.75), however, the pairwise correlations between the Nama and the other African populations are much weaker (∼0.79) than the pairwise correlations between the African populations (> 0.90).

**Figure 2:**
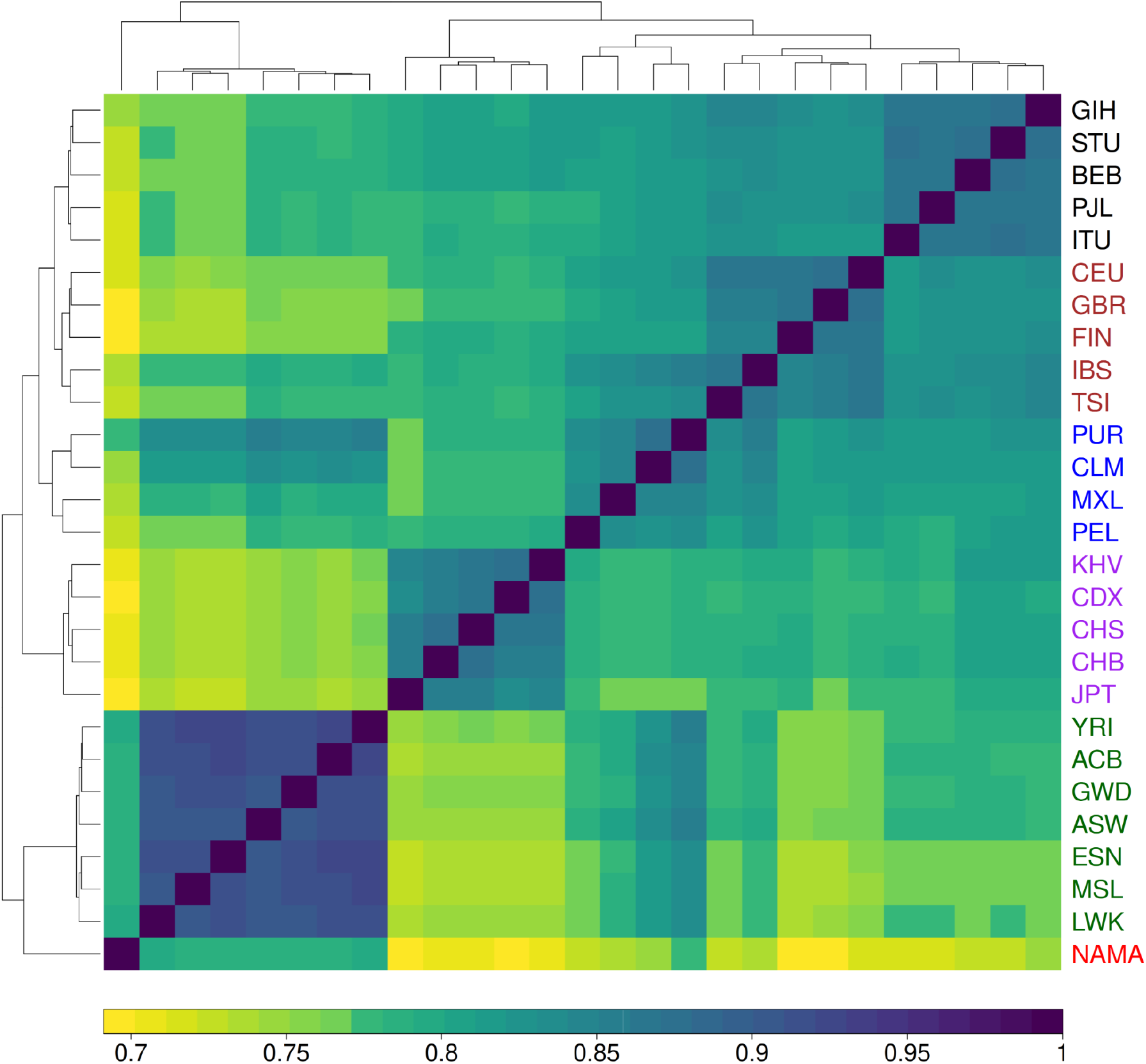
Heatmap indicating the Spearman rank correlation between the genetic maps of 27 populations, including the Nama, at a 2 kilobase resolution. The colour of the population labels represent distinct super-population groups, with the Nama highlighted in red. There is clear clustering according to super-population groups and the Nama recombination map correlates the best with other African populations.

The Spearman rank correlation mitigates potential differences in map length that would influence the Pearson correlation coefficient. Therefore, we neglect the magnitude of the recombination rate in favour of qualitative aspects of the maps. Inspecting the qualitative aspects of recombination maps is especially relevant when LD-based recombination maps are compared, since LD-based methods produce population recombination rates that need to be scaled using N_e_ and therefore assume an accurate estimate for N_e_.

### A Look at Fine-Scale Differences and Their Application in Selection Scans

The combined Phase II HapMap recombination map is derived from 270 individuals who represent four geographically diverse populations, including the Yoruba from Western Africa. It is sometimes used as a proxy (Vicente et al. 2019) for southern African populations, since all other available recombination maps derived from African populations are of western African ancestry, a globally diverse map is thought to be the best substitute. Even though population-specific recombination maps are similar at low resolutions, certain analyses, such as selection scans, might benefit from a high-resolution population-specific recombination map that accurately captures fine-scale differences. Figure 3 illustrates the recombination rate (cM/Mb) plotted over part of chromosome 1 for the combined Phase II HapMap recombination map (orange) and the inferred recombination map for the Nama (blue). The positions of regions of high recombination (hotspots) are largely concordant between the two maps and mainly differ in magnitude. However, in the region at 24.2 Mb there are hotspots present in the Nama recombination map that are absent from the combined Phase II HapMap recombination map. To further investigate the effects that these differences could have, we performed genome-wide selection scans on Nama SNP array data using the combined Phase II HapMap map and the Nama map. We focused on the integrated haplotype scores (iHS), a selection statistic which detects recent positive selection, by evaluating haplotype homozygosity for the ancestral and derived haplotypes extending from a locus of interest (Voight et al in 2006). iHS is most effective at detecting alleles that have been swept to intermediate frequencies, and it is among the most common statistics cited in other comparable selection scans in the Khoe-San.

**Figure 3:**
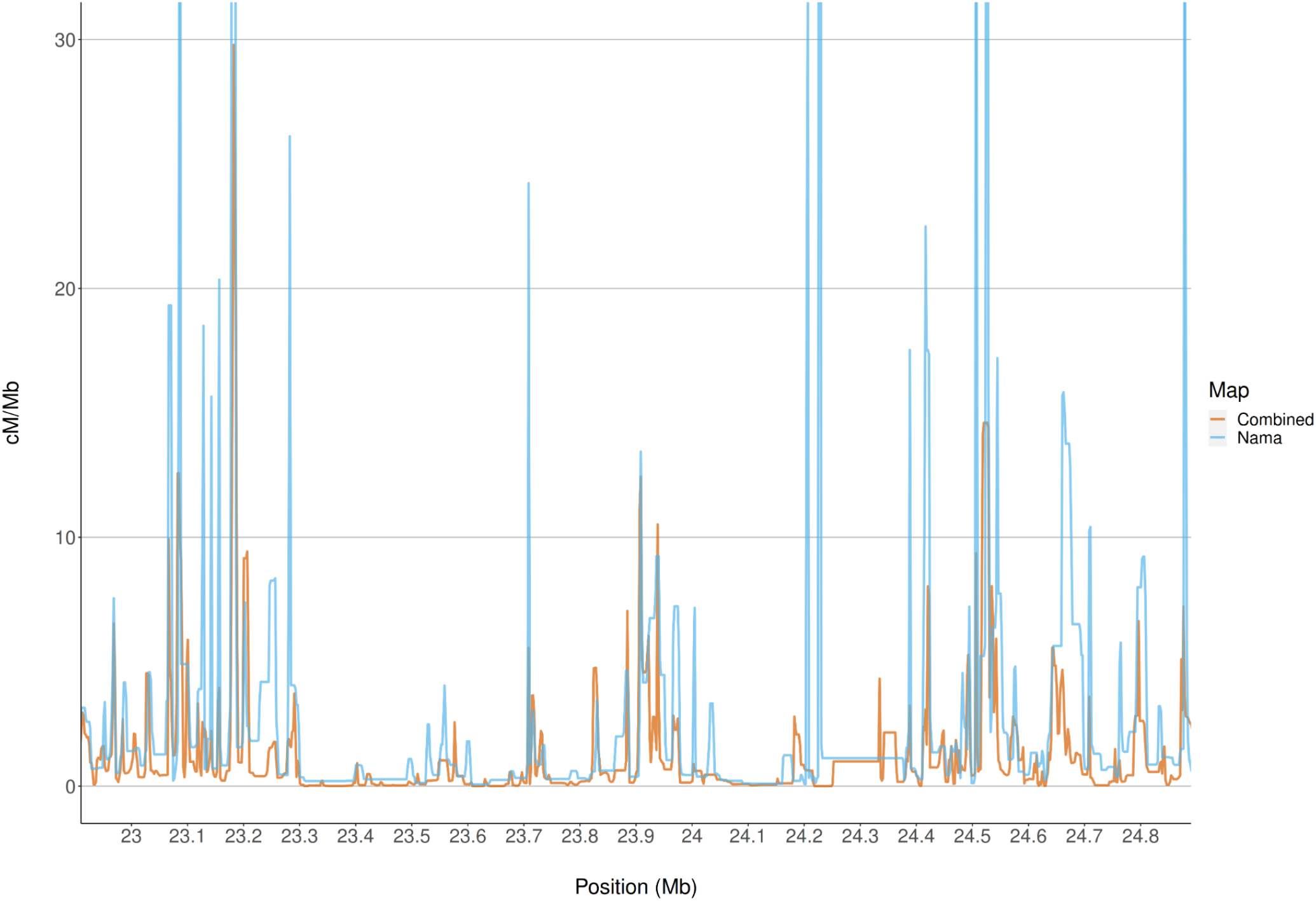
The recombination rate of the combined Phase II HapMap recombination map and the inferred recombination map for the Nama plotted over a segment of chromosome 1. There is a high degree of overlap between the maps across this region, but there are positions with recombination hotspots indicated by the Nama map that are not indicated by the combined Phase II HapMap map, e.g. at 24.2 Mb.

After taking the absolute value of the integrated Haplotype Scores (iHS) and filtering for the highest 1.0% of the scores, we found an overlap of 1504 candidate genes (50%) between the two maps. However, the run using the combined Phase II HapMap map and the run using the Nama map identified 808 and 713 unique candidate genes respectively (Fig. 4). The difference in the number of top 1.0% of hits is due to the change in the relative length of the maps. The Pearson correlation (r) between the iHS scores found using the combined Phase II HapMap and Nama maps is 0.93. We compiled a list of 131 candidate genes (Henn et al. 2011; Schlebusch et al. 2012; Vicente et al. 2019), previously identified using iHS, that are under selection in the Khoe-San and compared this list to our results. We found an overlap of three genes (*CTNNAL1, ALDH1A2* and *SYT14*) between the previously identified genes and the run using the combined Phase II HapMap map but only an overlap of one gene (*TRIM39*) between the previously identified genes and the run using the Nama map. *TRIM39* encodes for a ring finger protein associated with diseases including Behcet Syndrome; it regulates p21 and plays an important role in determining cell fate (Zhang et al 2012). Previous research has demonstrated that selection statistics such as iHS are sensitive to phasing, sample size and ascertainment bias (Granka et al. 2012). Our results indicate that a population-specific recombination map should also be considered in attempts to fine-map adaptive haplotypes.

**Figure 4:**
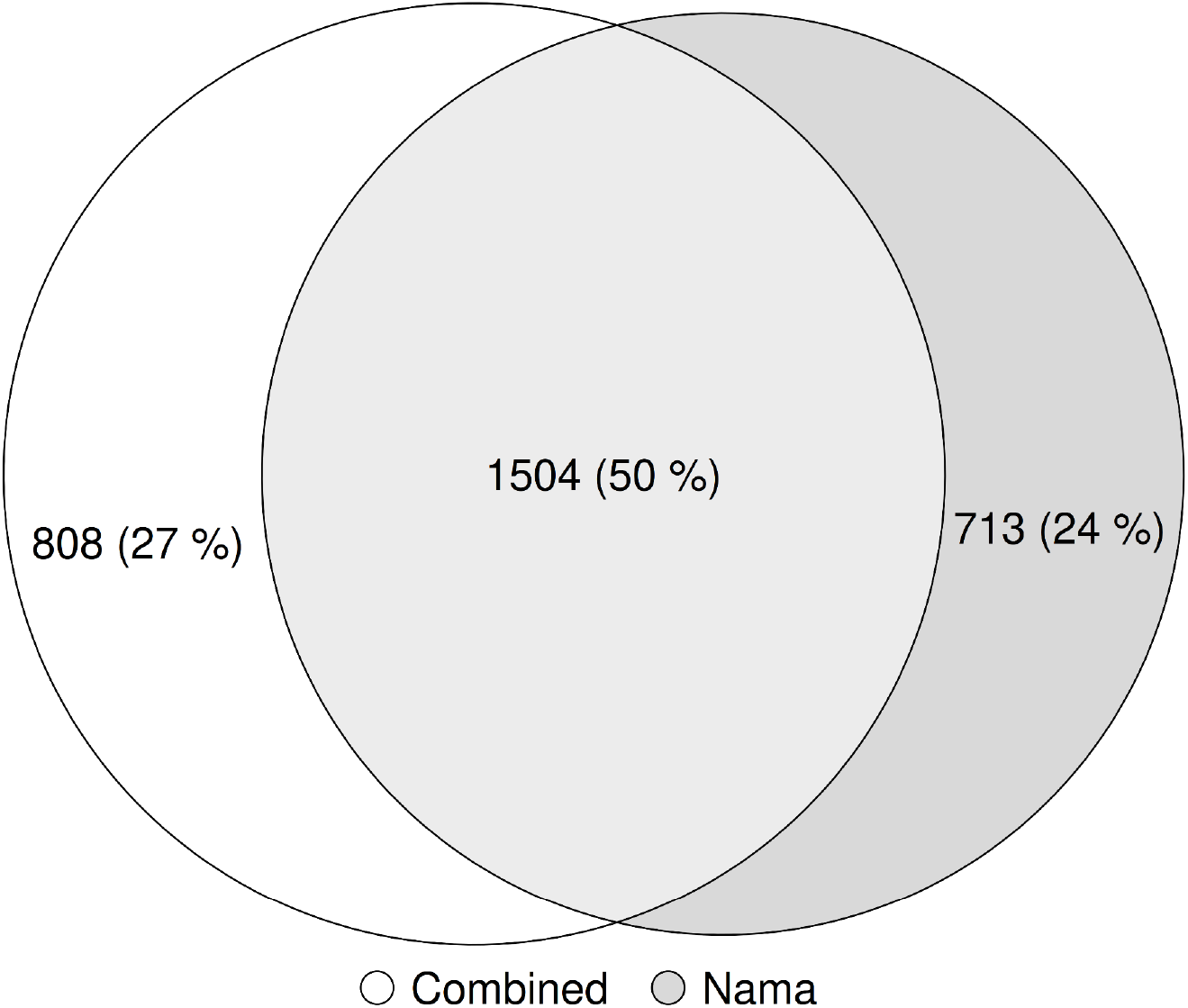
Venn diagram of the candidate genes found using the 1.0% highest selection scan results (absolute value iHS) for the selection scan using the combined Phase II HapMap map (white) and the selection scan using the Nama map (grey).

## Discussion

Recombination maps are important resources for epidemiological and evolutionary analyses, however, there are currently no recombination maps that represent southern African populations (Swart et al. 2020). The Nama, a southern African indigenous population, would likely produce a distinct recombination landscape from publicly available recombination maps, because of their complex demographic history. The recent rapid population decline (shown in Fig. 1) partially illustrates this complex history. Despite gene flow from Eastern African pastoralists ∼2000 years ago and recent admixture with Europeans, the Nama do not cluster with any of the continental groups that we have representative recombination maps for (Fig. 2). Therefore, their recombination landscape is indeed unique and epidemiological studies that involve the Nama or any other related populations, like other Khoe-San populations or southern African Bantu-speaking groups, would benefit from our inferred map. This recombination map also represents a likely upper bound on the extent of divergence we expect to see for a recombination map in humans and would be of interest to any researcher that wants to test the sensitivity of population genetic or GWAS analysis to recombination map input. Fine-scale differences in recombination can meaningfully alter the results of a selection scan (demonstrated in Figs. 3 and 4). However, it should be noted that recent studies found that population-specific recombination maps have little effect on phasing (Hassan et al. 2020), imputation (Hassan et al. 2020) and local ancestry inference (van Eeden, Uren, van der Spuy, et al. 2021). Therefore, the combined Phase II HapMap recombination map’s proxy status with regards to the Nama is dependent on the analysis that the map is used for.

Since it was not feasible to collect extended pedigree information and specimens and because of the demographic history of the Nama, it was evident that there would be numerous challenges to overcome in pursuing a novel population-specific recombination map that would be accurate and representative enough to be useful in epidemiological studies. There are many available techniques (van Eeden, Uren, Möller, et al. 2021) to infer the recombination rate and some have contrasting limitations which means that not all techniques would allow accurate, fine-scale estimates for a given dataset. Assuming limitless resources, we would have preferred pedigree-based methods, because these allow sex-specific recombination rate inference and rely on inferring individual recombination events between successive generations based largely on observed meioses. However, pedigree-based methods require many thousands of individuals to produce fine-scale maps (Halldorsson et al. 2019). Other options are IBD-based and LAI-based methods, but they too require in the order of a couple thousand individuals for fine-scale estimates (Zhou et al. 2020). Our small sample size (54 unrelated individuals) made LD-based methods the obvious choice for fine-scale estimates. However, there are many assumptions that accompany LD-based methods that make them less than ideal, for instance the assumption of a constant N_e_. Therefore, the complex demographic history of the Nama made demography-aware methods, like pyrho, the ideal compromise between data availability and accuracy. Even so, the population-specific recombination map presented here is likely an accurate representation of the recombination landscape of the Nama and future epidemiological and evolutionary research will benefit from this resource.

## Materials and Methods

### Inferring Demographic History

It has been shown that demographic history, especially recent bottlenecks, can greatly impact LD-based recombination inference. We, therefore, inferred the demographic history of the Nama to improve our recombination rate estimates. Two methods, SMC++ (v1.15.2) (Terhorst et al. 2017) and IBDNe (v23Apr20) (Browning and Browning 2015), were used and the results combined. See Figure 5 for an overview of the methods.

**Figure 5:**
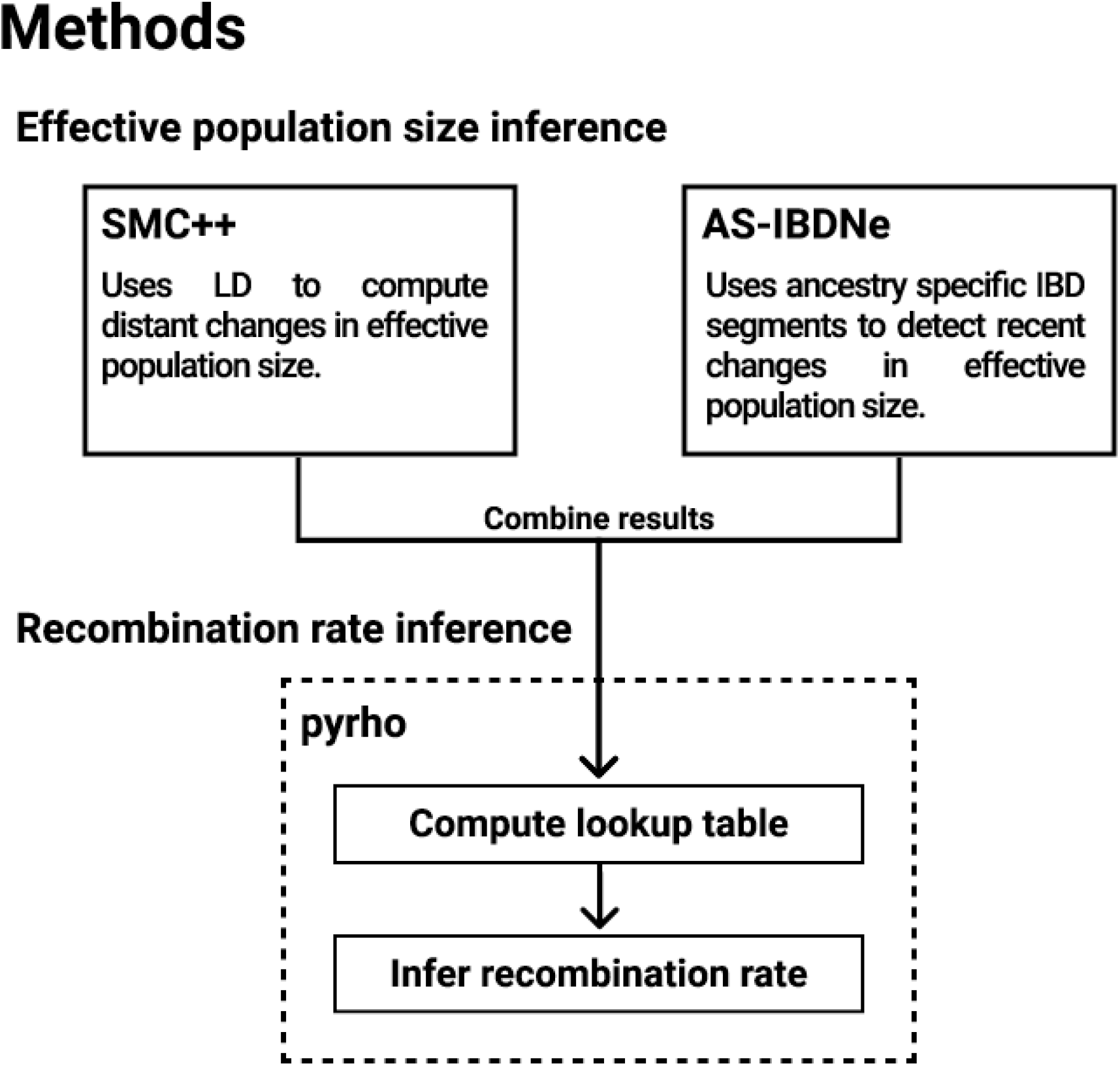
A brief overview of the methods.

SMC++ uses LD information to infer demographic histories and can infer divergence times between 6-120 kya with low error (Terhorst et al. 2017). A whole genome sequencing (WGS) dataset (EGAD00001006198) of 84 Nama individuals (54 unrelated) was used. The input for SMC++ was created separately for each chromosome, from the unrelated individuals in the WGS dataset, by using the *vcf2smc* program with 10 randomly selected “distinguished” (See Terhorst et al. (2017) for more information on this) individuals. The result is 10 separate datasets for each chromosome. This creates a composite likelihood which, according to the authors, may lead to improved estimates. A per-generation mutation rate of 1.25e-8 was assumed and all of the input files were then included in an estimate of the N_e_ through time using the *estimate* program. Since SMC++ regards uncalled regions as long runs of homozygosity, Stephen Schiffels’ mappability mask (created for human genome build GRCh37 using SNPable (Li 2009)) was used to mask regions of low mappability. All other default parameters were used.

IBDNe can infer the N_e_ size 4-50 generations into the past by using identity by descent (IBD) information. By separating IBD segments by ancestry before inferring the N_e_, one can obtain an estimate of N_e_ localized to each population ancestry. We developed a Snakemake pipeline (Fig. 6), called AS-IBDNe (https://github.com/hennlab/AS-IBDNe), to estimate ancestry specific N_e_ from a given SNP array dataset. The pipeline was adapted from the procedure used in Browning et al. (2018). We ran it on 84 other Nama individuals genotyped on the Multi-Ethnic Global Array (MEGA) (Martin et al. 2017). The pipeline takes in SNP-array data in plink binary file format, uses plink v1.9 (Chang et al. 2015) to break the data by chromosome, and shapeIT v2 (O’Connell et al. 2014) to phase the chromosomes. The dataset is then converted to VCF format using SHAPEIT2, and split into one file containing the reference individuals and one file containing the admixed individuals using BCFtools (Danecek et al. 2021). Next, RFMix v2.0 (Maples et al. 2013) is run on these two vcf files to estimate the ancestry of arbitrarily sized segments across the genome. Simultaneously, RefinedIBD and merge-ibd-segments.17Jan20.102.jar (Browning and Browning 2013) is run on the phased data to infer ibd segments and remove any gaps between them. The ancestries produced from RFMix v2.0 are then assigned to each IBD segment using a custom python script. This information is provided to the program IBDNe (Browning and Browning 2015), which produces estimates of historical population size for each ancestry. All the default parameters of RFMix, RefinedIBD and IBDNe were used except RFMix, which was run with 3 expectation maximization iterations and the *reanalyse-reference* flag, and IBDNe, which was run with the *mincM* flag set to 3. The combined Phase II HapMap recombination map was used whenever a recombination map was required during the inference. The output of SMC++ can be converted to a csv where the timescale and the N_e_ estimates are linear. The output from AS-IBDNe can then be added to the linear output from SMC++ and this file can then be used during recombination rate inference in pyrho (Spence and Song 2019).

**Figure 6:**
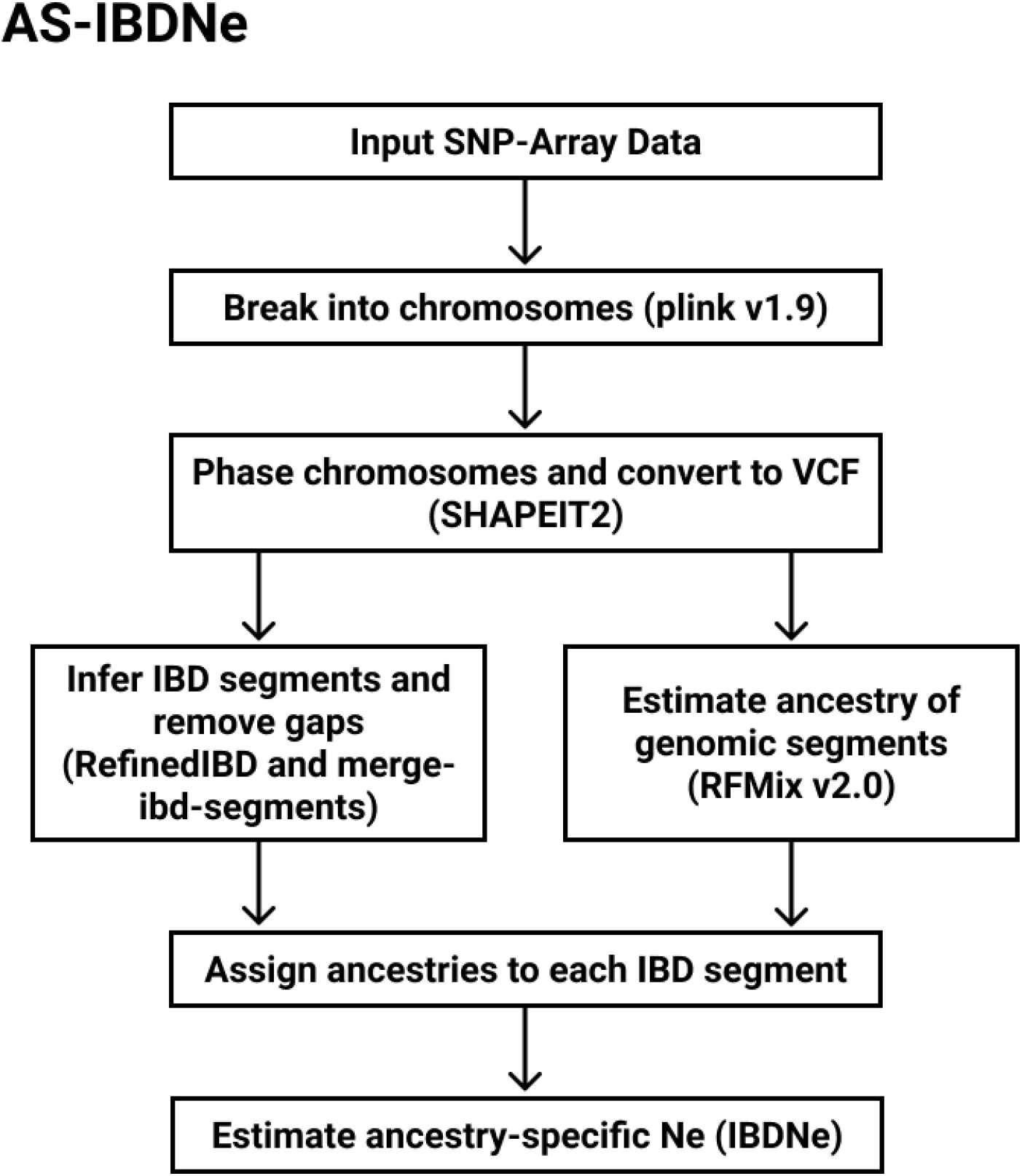
An overview of the AS-IBDNe pipeline.

### Recombination Rate Inference

Previous published guidelines (van Eeden, Uren, Möller, et al. 2021) aided the choice of recombination rate inference method and pyrho, a demography-aware LD-based method, was selected. We assumed the same per-generation mutation rate of 1.25e-8 for all the inference steps. The most computationally laborious task when using pyrho is the generation of a lookup table which enables subsequent processes to be computationally faster. The combined SMC++ / AS-IBDNe demographic history was used to generate a lookup table for the unrelated subset of 54 Nama WGS individuals using pyrho *make_table*. A convenient feature of this lookup table is that it is compatible with other recombination rate inference software, e.g LDhat (Auton and McVean 2007), which make use of exact two-locus sampling probabilities with the added benefit of already taking the specified demographic history into account. This lookup table and the combined demographic history were employed to find optimal hyperparameters to be used for recombination rate inference with pyrho *hyperparam*. The parameters that yielded the highest overall accuracy were a smoothness penalty of 15 and a window size of 30. These parameters and the lookup table were then used to infer the recombination rate with pyrho *optimise*. The output provides the per base pair per generation recombination rate for a given interval.

### Selection Scans

For the selection scans we used data from 104 Nama individuals who were genotyped on the Illumina Omni2.5 array as part of the African Genome Diversity Project. Close relatives were identified from demographic interviews and verified via allele-based kinship coefficients in Plink. Individuals with more than 50% European, and Damara or Herero admixture were excluded. Ancestry estimates were obtained using ADMIXTURE with k=6 possible ancestral clusters: Nama, Northern San, Near Eastern, East African Nilotic, West African and European (Lin et al. 2018). After QC, kinship and ancestry exclusions, we analyzed n=55 individuals. We calculated iHS using selscan 1.3.0 (Szpiech and Hernandez 2014) and default parameters. For the --map flag we used recombination rates from the custom Nama map in one run and from the combined Phase II HapMap in a second run. We filtered for the most extreme iHS scores (absolute value) by taking the highest 1.0% of the scores.

We annotated these positions using the gene range list provided by Plink ((https://www.cog-genomics.org/plink/1.9/resources). We compared the candidate genes found in each run of selscan to create a Venn Diagram. We also calculated the Pearson correlation between iHS scores for each SNP as calculated by each run of selscan.

## Data Access

Sequence data has been deposited at the European Genome-phenome Archive (EGA), which is hosted by the EBI and the CRG, under accession number EGAD00001006198. The recombination map inferred for the Nama can be found at https://github.com/TBHostGen/nama-recombination-map.

## Acknowledgements

This research was funded (partially or fully) by the South African government through the South African Medical Research Council and the National Research Foundation. GvE was supported by the DSI-NRF Innovation Doctoral Scholarship. This research was supported by NIH grant R35GM133531 (to BMH). The content is solely the responsibility of the authors and does not necessarily represent the official views of the National Institutes of Health. The authors would also like to thank Prof. Carina Schlebusch and Dr. Torsten Günther for providing previously published data on the demographic history of the Nama.

